# Whole genome analyses reveal weak signatures of population structure and environmentally associated local adaptation in an important North American pollinator, the bumble bee *Bombus vosnesenskii*

**DOI:** 10.1101/2023.03.06.531366

**Authors:** Sam D. Heraghty, Jason M. Jackson, Jeffrey D. Lozier

**Author notes:** Corresponding authors: Sam D. Heraghty Mailing address: Box 870344, University of Alabama, Tuscaloosa, AL, 35487.

## Abstract

Studies of species that experience environmental heterogeneity across their distributions have become an important tool for understanding mechanisms of adaptation and predicting responses to climate change. We examine population structure, demographic history, and environmentally associated genomic variation in *Bombus vosnesenskii*, a common bumble bee in the western U.S.A., using whole genome resequencing of populations distributed across a broad range of latitudes and elevations. We find that *B. vosnesenskii* exhibits minimal population structure and weak isolation by distance, confirming results from previous studies using other molecular marker types. Similarly, demographic analyses with Sequentially Markovian Coalescent (SMC) models suggest that minimal population structure may have persisted since the last interglacial period, with genomes from different parts of the species range showing similar historical effective population size (*N*_e_) trajectories and relatively small fluctuations through time. Redundancy analysis revealed a small amount of genomic variation explained by bioclimatic variables, and environmental association analysis with latent factor mixed modeling (LFMM2) identified few outlier loci that were sparsely distributed throughout the genome. Some outlier loci were in genes with known regulatory relationships, suggesting the possibility of weak selection, although compared to other species examined with similar approaches, evidence for extensive local adaptation signatures in the genome was relatively weak. Overall, results indicate *Bombus vosnesenskii* is an example of a generalist with a high degree of flexibility in its environmental requirements that may ultimately benefit the species under periods of climate change.

## 1 Introduction

Studying species that experience environmental heterogeneity is a key component in understanding phenomena like local adaptation and predicting species responses to climate change (Hoffmann, Weeks, & Sgro, 2021; Savolainen, Lascoux, & Merilä, 2013; Sears, Riddell, Rusch, & Angilletta, 2019). The degree of spatial and environmental heterogeneity across a species range can have important effects on gene flow and population structure (Y. Li et al., 2017; Manel, Schwartz, Luikart, & Taberlet, 2003). Spatially complex landscapes have features that might constrain gene flow, increasing population genetic structure of inhabitants compared to more homogeneous landscapes (Rahbek et al., 2019; Wang & Singh, 2019) and also harbor substantial abiotic variation that could drive local adaptation in populations from dissimilar environments (Andrews et al., 2022; Antoniou et al., 2023; Capblancq, Fitzpatrick, Bay, Exposito-Alonso, & Keller, 2020; Heraghty, Rahman, Jackson, & Lozier, 2022). Genomic data can be used to understand current and historical population structure as well as genome-environment associations that might signal local adaptation and can thus provide multiple types of information about evolutionary responses to complex landscapes in widespread species.

Although natural selection is expected to shape genetic variation in species distributed over large biogeographic gradients, not all species experience and respond to spatial or environmental heterogeneity in the same way (Gallegos, Hodgins, & Monro, 2023; Hartke et al., 2021; J. M. Jackson et al., 2020). For instance, species capable of long-distance dispersal or characterized by generalist individual phenotypes across a range of environmental conditions might show weak signals of local adaptation (Lenormand, 2002; Räsänen & Hendry, 2008). This phenomenon has been observed in a variety of taxa including marine invertebrates (Lal, Southgate, Jerry, Bosserelle, & Zenger, 2016), butterflies (Melero et al., 2022), and trees (De la Torre, Wilhite, Neale, & Slotte, 2019). Landscape genomics provides a lens for assessing the challenges of a heterogenous environment by revealing specific environmental features that drive population structure (Capblancq & Forester, 2021), and has also become a widely used tool for detecting signals of environmental adaptation (Capblancq, Morin, Gueguen, Renaud, & Bazin, 2020; De la Torre et al., 2019; Heraghty et al., 2022; J. M. Jackson et al., 2020), especially as whole genome data becomes more easily obtained.

Demographic responses to past climatic fluctuations can provide another important dimension for understanding environmentally associated genomic variation. In widespread species, for example, distinct demographic histories (e.g. bottlenecks) across a distribution can influence on how populations adapt to their contemporary environment, and testing for parallel or dissimilar effective population size (*N*_e_) trajectories through time can provide context for inferences of contemporary population structure or local adaptation (Ahrens et al., 2018). A common tool for inferring *N_e_* is a class of methods known as Sequential Markovian Coalescence (SMC) models (Lozier, Strange, & Heraghty, 2023; Mather, Traves, & Ho, 2020; Nadachowska-Brzyska, Burri, Smeds, & Ellegren, 2016), which can provide estimates of temporal variation in *N_e_* using genomes from single individuals or populations. SMC approaches can reveal how populations have changed in size over tens of thousands of years, and thus reveal whether past climate conditions may have affected genetic variation in similar or distinct ways across a species range (Lozier et al., 2023; Taylor et al., 2021). Such information may also be helpful in interpreting contemporary patterns of environmental association since complex histories could constrain genetic diversity which ultimately may limit the ability of organisms to adapt to their environment (Reed & Frankham, 2003).

In this study, we employ a whole genome resequencing (WGR) approach to evaluate population structure, demographic history, and potential targets of adaptation in a common bumble bee, *Bombus vosnesenskii* Radowski. *Bombus vosnesenskii* is one of the most common bumble bee species in the westernmost parts of North America, with a range extending from southern California, USA through British Columbia, Canada (Cameron et al., 2011; Fraser, Copley, Elle, & Cannings, 2012; J. Koch, Strange, & Williams, 2012; Stephen, 1957). The latitudinal range of *B. vosnesenskii* indicates that the species is capable of inhabiting a broad environmental niche, as well as occurring at a range of elevations from sea level to 2,400 m in elevation (Stephen, 1957). Like many bumble bees, *B. vosnesenskii* provides important pollination services (Greenleaf & Kremen, 2006; Strange, 2015; Velthuis & Doorn, 2006) and is the only native western US bumble bee commercially available for pollination. Thus, in addition to better revealing patterns of evolutionary adaptation to heterogeneous environments in widespread species, improved understanding of regional genomic variation in *B. vosnesenskii* could be of value for breeding or assessing risks associated with bees used in commercial contexts (Lozier & Zayed, 2016), and for understanding potential risks to pollination from range shifts under climate change (H. M. Jackson et al., 2022; Kerr et al., 2012).

*Bombus vosnesenskii* is among the most well-studied bumble bees in the United States in terms of its geographic range, habitat use, physiology, and ecology (Heinrich & Kammer, 1973; J. Koch et al., 2012; Mola, Miller, O’Rourke, & Williams, 2020a; Pimsler et al., 2020; Stephen, 1957), although no study to date has investigated the species using range-wide whole genome resequencing. Prior information leads to several hypotheses relating to the effects of landscape heterogeneity on genomic variation in *B. vosnesenskii*. Microsatellite and reduced representation (RADseq) single nucleotide polymorphism (SNP) datasets have largely found evidence of minimal population structure and overall genetic homogeneity at the continental scale (J. M. Jackson et al., 2018; S. Jha, 2015; Lozier, Strange, Stewart, & Cameron, 2011). RADseq also revealed relatively few outlier loci associated with bioclimatic or elevational variables compared to a related narrower-niche montane species, *B. vancouverensis* (J. M. Jackson et al., 2020).

Foraging in *B. vosnesenskii* appears uninhibited by certain complex landscapes like forests (Mola et al., 2020a) and can benefit from pulses in floral availability associated with wildfire disturbance (Mola, Miller, O’Rourke, & Williams, 2020b), potentially indicating that movement of reproductive castes, and thus gene flow, could also be unimpeded or facilitated by landscape heterogeneity. Consistent with the hypothesis of *B. vosnesenskii* genetic homogeneity, range-wide morphological analyses have also found little evidence of variation in functional traits like body size or wing loading across latitude or altitude compared to *B. vancouverensis* (Lozier et al., 2021). There is evidence, however, that some spatial or environmental landscape features can impact *B. vosnesenskii* demography by influencing nesting density (Shalene Jha & Kremen, 2013a) and introducing local or regional barriers to gene flow (J. M. Jackson et al., 2018; S. Jha, 2015; Shalene Jha & Kremen, 2013b). Physiological assays have also detected variation in critical thermal minima (CT_min_) across latitudes and elevation that could indicate adaptive variation associated with local cold temperatures (Pimsler et al., 2020). Overall, *B. vosnesenskii* appears well-suited to occupying heterogenous landscapes, but with some potential for environmental features to shape aspects of the species’ evolution that may be clarified using the large genetic marker sets from whole genomes.

Here, we generate WGR data from *B. vosnesenskii* workers sampled from diverse environments across California and Oregon to test several hypotheses relating to population structure and local adaptation. WGR data is a powerful tool for detecting adaptation and can overcome potential shortcomings of previously used approaches such as reduced representation sequencing (e.g., RADseq) (Jackson et al. 2020), which may not detect all possible targets of selection when linkage blocks are small (Fuentes-Pardo & Ruzzante, 2017), as in bumble bees (Stolle et al., 2011). Whole genome data enables analyses that infer species’ demographic histories which can complement contemporary landscape genomics (Beichman, Huerta-Sanchez, & Lohmueller, 2018; de Greef et al., 2022; Iannucci et al., 2021). First, based on prior results of near-panmixia from other genetic markers, we examine population structure and test for any influence of spatial-environmental variation that may not have been apparent in earlier studies.

We also use SMC methods to test whether patterns of genetic structure or homogeneity detected in current populations is likewise reflected by divergent or parallel patterns of historical effective population size trajectories across the *B. vosnesenskii* range. Last, we expand upon previous RADseq data by performing genome scans at the fine resolution afforded by WGR data to identify potential signals of environmental adaptation across the *B. vosnesenskii* range.

## 2 Materials and Methods

### 2.1 Sample Collection, DNA extraction and Sequencing

*Bombus vosnesenskii* workers (diploid females) were selected for whole genome resequencing from previously collected samples (J. M. Jackson et al., 2018, 2020) that are representative of elevational extremes sampled across a range of latitudes in California and Oregon (36.5°N – 45.3°N latitude and 49m – 2797m above sea level) (Fig 1, Table 1). Our sampling represents a series of six relatively low and high elevation site-pairs nested across a range of latitudes, which should cover the breadth of environmental conditions experienced by *B. vosnesenkii,* such as mean annual temperature from 3°C- 17°C and annual precipitation from 369 mm – 2,177 mm (from WorldClim v2 (Fick & Hijmans, 2017). Briefly, samples were collected at each site via sweep netting, placed on ice for identification (being especially careful to exclude the phenotypically similar *B. caliginosus*), and then placed in 100% ethanol on dry ice, before final storage in ethanol at -80°C. Based on prior estimation of relatedness with RADseq data, selected workers should represent independent colonies. See Jackson et al. 2018 for a more complete description of sampling and the study region.

**Figure 1:**
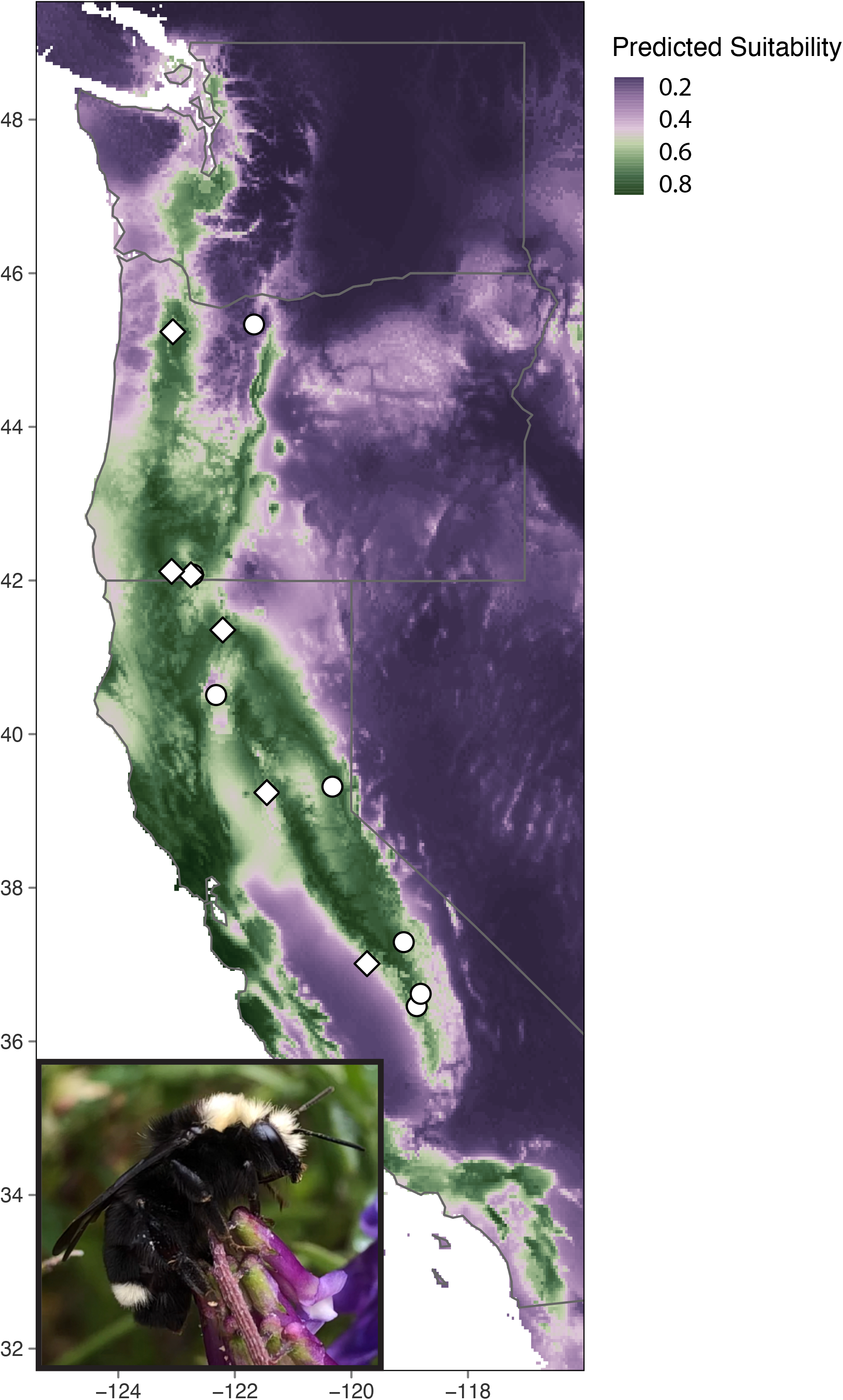
Map showing Maxent (Phillips, Anderson, Dudík, Schapire, & Blair, 2017) range of the *B vosnesenskii* using select environmental variables (BIO1, BIO3, and BIO12) with presence absence data from (Cameron et al., 2011). White diamonds indicate sites that were used for PSMC analysis.

**Table 1:** Summary of sampling sites and average nucleotide diversity (π) per population. * OR08.2012 and OR09.2012 were pooled together as a single population when calculating nucleotide diversity.

| Site | Number of samples | Latitude | Longitude | Elevation<br>(m) | $\pi$ |
| --- | --- | --- | --- | --- | --- |
| CA01.2015 | 8 | 36.458 | -118.879 | 313 | 0.2745 |
| CA04.2012 | 11 | 41.356 | -122.208 | 2261 | 0.2743 |
| CA06.2015 | 11 | 37.012 | -119.732 | 131 | 0.27478 |
| CA11.2013 | 12 | 39.316 | -120.326 | 2164 | 0.2745 |
| CA11.2015 | 7 | 39.237 | -121.451 | 49 | 0.2747 |
| CA12.2015 | 10 | 40.508 | -122.322 | 138 | 0.2746 |
| CA13.2015 | 8 | 36.619 | -118.811 | 2268 | 0.2751 |
| CA29.2015 | 9 | 37.291 | -119.102 | 2796 | 0.2746 |
| OR02.2015 | 5 | 45.238 | -123.063 | 53 | 0.2741 |
| OR05.2016 | 10 | 45.333 | -121.670 | 1698 | 0.2743 |
| OR08.2012 | 2 | 42.076 | -122.717 | 2055 | 0.2747* |
| OR09.2012 | 8 | 42.073 | -122.754 | 2134 | 0.2747* |
| OR11.2012 | 10 | 42.118 | -123.085 | 518 | 0.2746 |

DNA was extracted using the Qiagen DNeasy Blood and Tissue kit (Hilden, N.R.W., Germany). Some samples were sequenced using whole genome shotgun libraries prepared via the NEBnext Ultra II FS DNA kit (Ipswich, MA, USA) with subsequent 150 bp paired-end sequencing using Illumina NovaSeq 6000 technology (Psomagen, Rockville MD). The remaining sample libraries were prepared by the Genomics and Cell Characterization Core Facility at the University of Oregon and sequenced across three separate lanes of an Illumina Hiseq 4000 instrument.

### 2.2 Read mapping, filtering, and variant calling

Sequencing reads were processed with bbduk v37.32 (Bushnell 2020) to remove adaptors, trim low quality bases, and remove short reads (ktrim=r k=23 mink=11 hdist=1 tpe tbo ftm=5 qtrim=rl trimq=10 minlen=25). Read quality was evaluated using FastQC v0.11.5 (Andrews 2010) before being mapped to the *B. vosnesenskii* reference genome (NCBI RefSeq ID: GCF_011952255.1) (Heraghty et al., 2020) using BWA mem v0.7.15-r1140 (H. Li & Durbin, 2009). Samtools v1.10 (H. Li et al., 2009) was used to convert the SAM files to BAM files and Picard tools v2.20.4 (Broad Institute 2019) was used to sort, mark duplicates, and index the BAM files. Single nucleotide polymorphisms (SNPs) were called using freebayes v1.3.2 (Garrison & Marth, 2012) and filtered following Heraghty et al. (2022). An initial round of filtering was performed to remove low-quality variants using vcftools v0.1.13 (Danecek et al., 2011) with the following flags: --remove-indels --min-alleles 2 --max-alleles 2 --minQ 20 -- minDP 4 --max-missing 0.75. Subsequent filtering was designed to remove SNPs with unusually high coverage (> 2x average coverage), excess heterozygosity (--hardy flag in vcftools), located on small scaffolds (< 100 kb in size), and with a minor allele frequency (MAF) ≤0.05 to filter SNPs that might have arouse from repeat regions or sequencing artefacts and limit the effects of low frequency variants. A final round of filtering was done to remove differences that might arise from using data generated on different sequencing platforms, as recommended by (De-kayne et al., 2021), using a higher SNP quality filter (min Q of 30 and min GQ of 20).

### 2.3 Environmental Variable Selection and Population Structure

Environmental variables (19 Bioclim variables) at a 0.5 arcminute resolution were obtained from WorldClim v2 (Fick & Hijmans, 2017). To reduce correlation between variables we used an item clustering analysis (*iclust* function with default settings) from the *psych* v.2.1.9 package (Revelle 2020) in R v 4.2.0 (R Core Team 2022) and retained a single variable per correlated variable cluster. Retained variables were: BIO1 – annual mean temperature, BIO3 – isothermality, and BIO12 – annual precipitation. Elevation was also included as a variable of interest despite being partly correlated with other environmental variables as it may capture unique environmental information that could be relevant for bumble bees such as air density and oxygen availability (Cheviron & Brumfield, 2012; Dillon, 2006; Heraghty et al., 2022).

To assess population structure, the fully filtered vcf file was converted into a genlight object via the *vcfR2genlight* function in the *vcfR* v1.12.0 package in R (Knaus & Grünwald, 2017). Global *F*_ST_ (Weir & Cockerham 1984 method (Weir & Cockerham, 1984)) was calculated using *SNPRelate* 1.30.1 (Zheng et al., 2012) in R. Pairwise *F*_ST_ was calculated between all sampling localities using the stamppFst function from the *dartR* v1.9.9.1 package in R (Gruber, Unmack, Berry, & Georges, 2018). A geographic distance matrix for all sampling points was created using the *distm* function from the *geosphere* v1.5-14 package in R using coordinates of each site. A Mantel test (statistical testing using 1000 permutations) was performed on pairwise *F*_ST_ and geographic distance matrices to test for signatures of isolation by distance (Slatkin, 1993) using the mantel function from the *vegan* v2.5-7 package in R (Oksanen et al 2020). To visualize population structure, the *gl.pcoa* function from the *dartR* v1.9.9.1 package in R (Gruber et al., 2018) was used to conduct a Pearson PCA. Nucleotide diversity (π) was calculated for each population using the – site-pi flag in vcftools v0.1.13 (Danecek et al., 2011).

We also evaluated the relative contribution of spatial and environmental factors on the partitioning of genetic variation in *B. vosnesenskii* using partial redundancy analysis (pRDA) (Capblancq & Forester, 2021). A full RDA model was created with *rda* function in the *vegan* v2.5-7 package in R using the selected environmental variables BIO1, BIO3, BIO12, elevation, and using latitude as a proxy for geographic effects. The pRDA model accounting for environmental effects (hereafter referred to as “Environmental pRDA”) used the following model (BIO1 + BIO3 + BIO12 + elevation | latitude) and the pRDA model accounting for geographic distance effects (hereafter referred to as “Geography pRDA”) used the following model (latitude | BIO1 + BIO3 + BIO12 + elevation). We also examined models with just environmental variables or latitude separately (i.e. without the partial effects), which we refer to as the “Environmental RDA” and “Geography RDA”, respectively. All models were assessed for significance using the *anova* function in R following the protocol from (Capblancq & Forester, 2021).

### 2.4 Demographic Inference

To test whether populations from different parts of the *B. vosnesenskii* range exhibit parallel or distinct recent evolutionary histories, we inferred historical *N_e_* trends using SMC analyses of whole genomes (H. Li & Durbin, 2011; Mather et al., 2020). SMC methods use the distribution of heterozygous and homozygous sites within loci to calculate coalescence rates that provide insight into past demographic events (Beichman et al., 2018; H. Li & Durbin, 2011).

This approach is a common tool used when WGR data are available, even for small numbers of individuals, as methods can make use of even single diploid genomes (Mather et al., 2020; Nadachowska-Brzyska et al., 2016). We used PSMC (H. Li & Durbin, 2011) for demographic analyses, which is the original but still widely employed (Lozier et al., 2023; Morin et al., 2021; Nadachowska-Brzyska et al., 2016; Patil et al., 2021; Skovrind et al., 2021) SMC method, and generally followed methods in Lozier et al. (2023). PSMC uses whole genome data from single individual diploid samples to estimate coalescence rates to track *N*_e_ history. Because SMC methods generally perform best with high coverage and long scaffolds (Mather et al., 2020), we only ran PSMC for *B. vosnesenskii* workers with ≥18x mean coverage (Nadachowska-Brzyska et al., 2016) and for reference genome scaffolds ≥500kb in length (Gower et al., 2018). Following methods described in the PSMC manual, the PSMC input files were constructed from the filtered BAM files using Samtools v1.10 mpileup with base and mapping quality set to 30. The consensus sequence was generated using bcftools call, converted to fastq format with Samtools vcfutils.pl vcf2fq, and converted to the psmcfa format for demographic inference using the psmc program. The default p parameter describing temporal intervals of “4+25*2+4+6" was used, which is typically useful for a range of taxa (Patil & Vijay, 2021). Results were plotted using the plot_psmc.py script provided with PSMC. To convert the results into real time, we used the direct mutation rate estimate of 3.6 × 10^-9^ per site per year from the bumble bee *Bombus terrestris* (Liu et al., 2017) and a generation time of 1 year.

### 2.5 Identifying environmentally associated genomic loci

Environmental Association Analysis (EAA) is often used to detect genetic variants with unusual allele frequency distributions that suggest a possible role in local adaption (e.g., “outlier” loci)(Ahrens et al., 2018). Here we use a common EAA method, latent factor mixed modeling with *LFMM2* from the *LEA* v3.0.0. R package (Gain & François, 2021), which uses a least-squares approach to identify SNPs with a significant association with a given variable while controlling for population structure. The likely number of population clusters (*k*) used for background population structure control was determined with the *sMNF* function from the *LEA* v3.0.0. R package (Gain & François, 2021), with *k* representing the value that had the smallest cross-entropy selected from a range of *k* = 1 – 10. To correct for multiple testing with the large number of genetic markers, raw *p-*values from LFMM2 were transformed into *q-*values using the *q-value* v2.20.0 R package (Storey et al. 2022). SNPs were considered significantly associated with a given variable at a threshold of *q ≤* 0.05.

## 3 Results

### 3.1 Data summary

A total of 111 individuals were sequenced to an average of 21,053,016 reads per individual and variant calling yielded an initial dataset of 22,105,684 SNPs. The final dataset consisted of 1,091,021 SNPs at a 5% minor allele frequency with an average coverage of 13.39x SNP^-1^ individual^-1^ and < 25% missing data SNP^-1^ (98.02% complete data across all samples).

### 3.2 Population structure and demographic history

Consistent with prior results in *B. vosnesenskii*, there was minimal population structure, with global *F*_ST_ = 0.001. The PCA also revealed little population structure and weak spatial clustering, but with a trend towards individual loadings related to the latitude at which the samples were collected (Fig 2a). This trend likely reflects the weak but significant isolation by distance at a range-wide scale (Mantel *r*: 0.374, *P =* 0.006; Fig 2b). The RDA models also supported minimal population structure, with all models explaining little variation, with the full model explaining only ∼5% of the variation (Table 2). Neither of the partial RDA models were significant, but all three RDA models without partial effects (Full model, Environment RDA, and Geography RDA) were significant. The Geography RDA indicates that most variation is likely attributable to weak isolation by distance as suggested by above analyses, with samples clustering together by approximate sampling latitude along the x-axis (Fig. 2), but with a significant amount of unexplained variation (captured by the y-axis).

**Figure 2:**
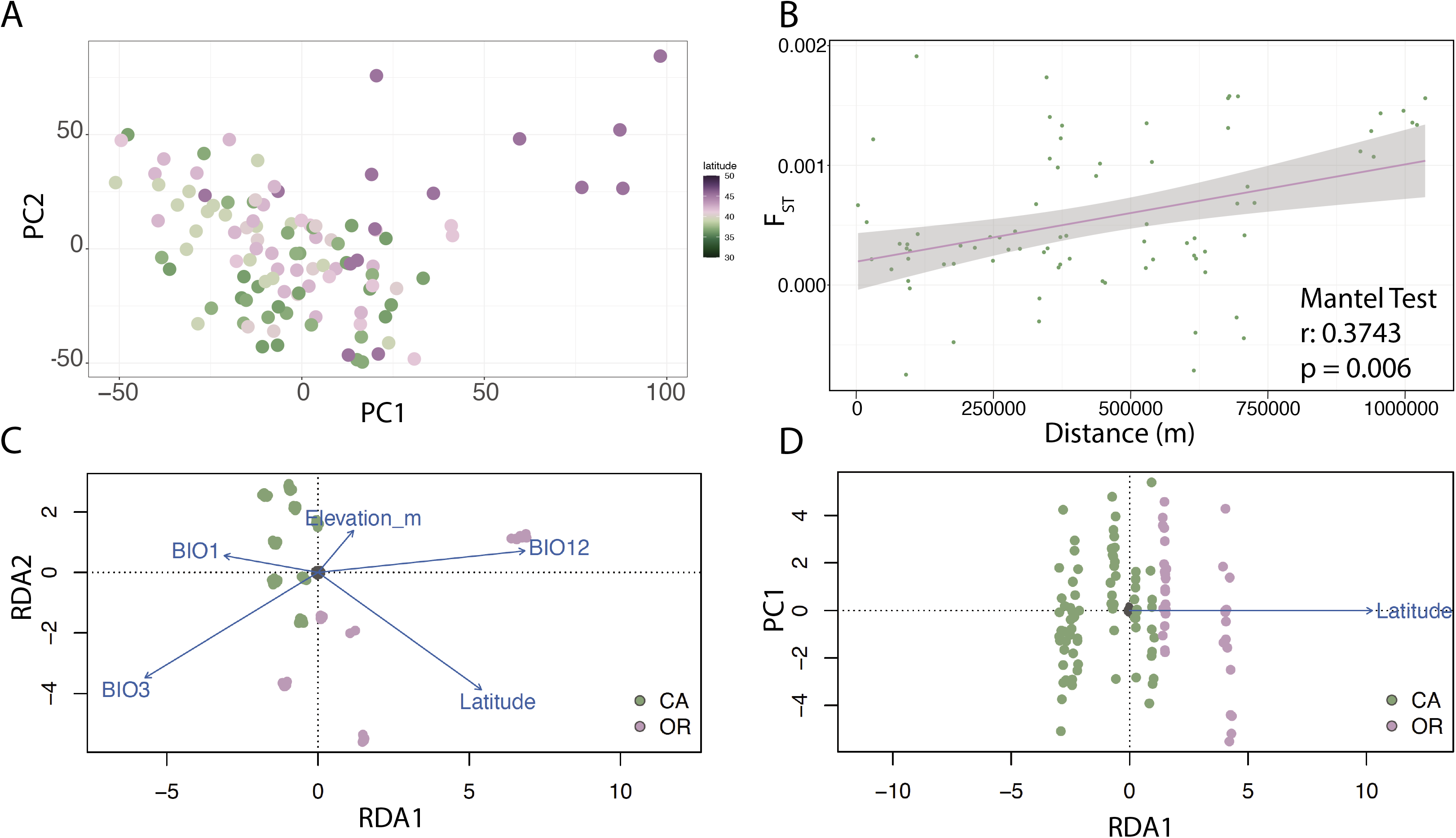
A) Pearson Principal Component analysis with samples colored by latitude of origin. B) Results of the isolation by distance analysis. C) The full RDA model containing all selected environmental variables as well as latitude (a proxy of distance) with samples colored by state of origin. Note this model is essentially identical to the RDA with only environmental variables, so we only present the results of the full model. D) The geography RDA model with samples colored by state of origin.

**Table 2:** A) Results of the pRDA analysis reporting the results of the full RDA model (BIO1 + BIO3 + BIO12 + Elevation + Latitude), the results of the Environment pRDA (BIO1 + BIO3 + BIO12 + Elevation | Latitude) and the results of the Latitude pRDA (Latitude | BIO1 + BIO3 + BIO12 + Elevation). It also shows that amount of confounded variation (variation that could be associated with either pRDA model), and the amount of unexplained variation. B) Results of different the full RDA model (BIO1 + BIO3 + BIO12 + Elevation + Latitude) (note this is the same as in table 1A), the results of the Environment RDA (BIO1 + BIO3 + BIO12 + Elevation) and the result of the Latitude RDA (Latitude).

| Model | Inertia | R<sup>2</sup> | *p* | Proportion of explainable variance | Proportion of total variance |
| --- | --- | --- | --- | --- | --- |
| Full (Env + Lat) | 13985 | 0.037 | 0.001 | 1 | 0.0486 |
| Environment pRDA | 11140 | 0.009 | 0.09 | 0.7966 | 0.0387 |
| Latitude pRDA | 2781 |  | 0.503 | 0.1989 | 0.0097 |
| Confounded | 64 |  |  | 0.0046 | 0.0002 |
| Unexplained | 273535 |  |  |  | 0.9514 |
| Total | 287520 |  |  |  | 1 |

Table 2: A) Results of the pRDA analysis reporting the results of the full RDA model (BIO1 + BIO3 + BIO12 + Elevation + Latitude), 847 the results of the Environment pRDA (BIO1 + BIO3 + BIO12 + Elevation | Latitude) and the results of the Latitude pRDA (Latitude | 848 BIO1 + BIO3 + BIO12 + Elevation).
| Model | Inertia | R<sup>2</sup> | *p* | Variance Explained | Total Variance |
| --- | --- | --- | --- | --- | --- |
| Full (Env + Lat) | 13985 | 0.037 | 0.001 | 0.0486 | 287520 |
| Environment RDA | 11205 | 0.009 | 0.001 | 0.0386 | 290301 |
| Geography RDA | 2845 |  | 0.001 | 0.0095 | 298660 |

**Table 3:**
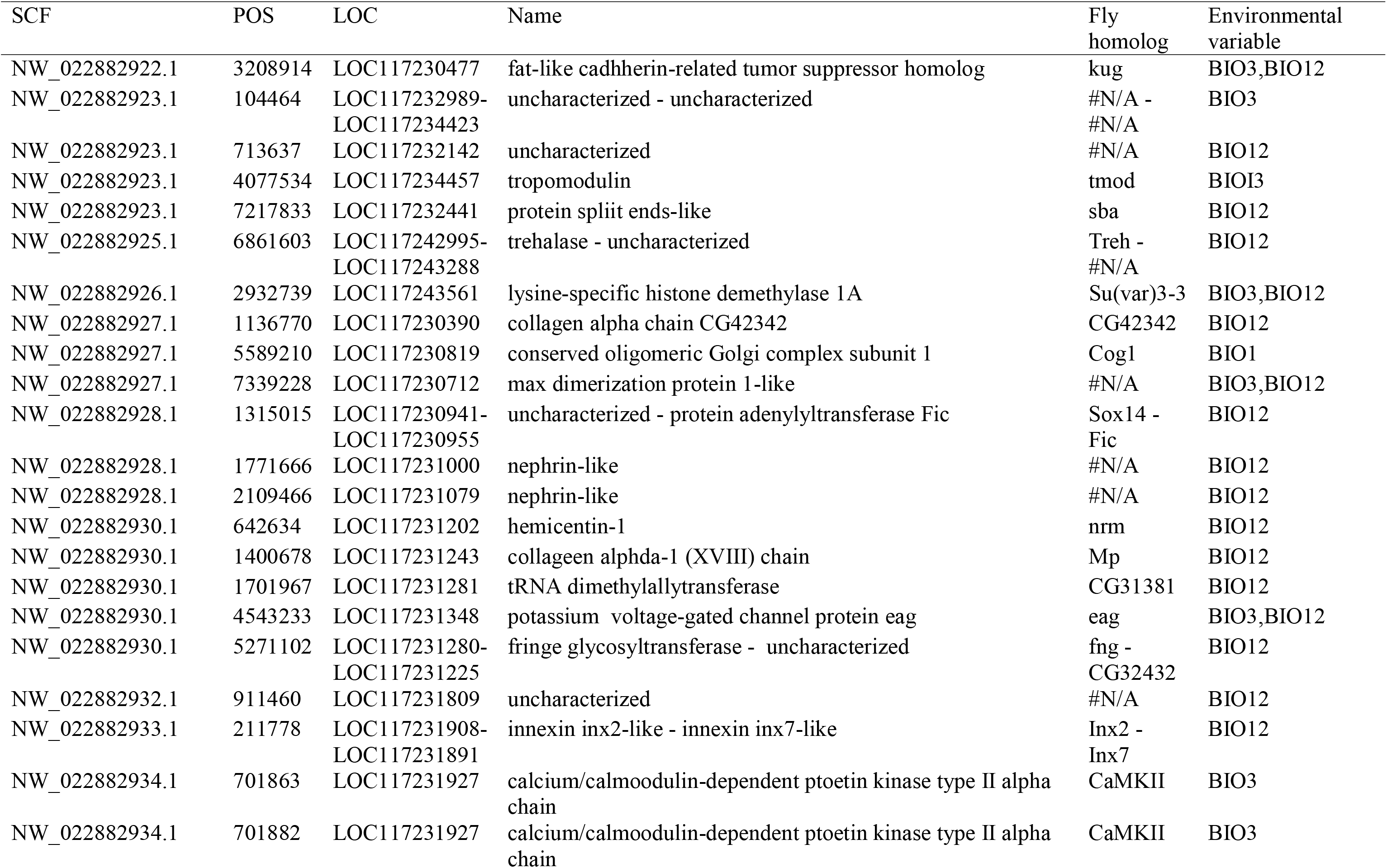
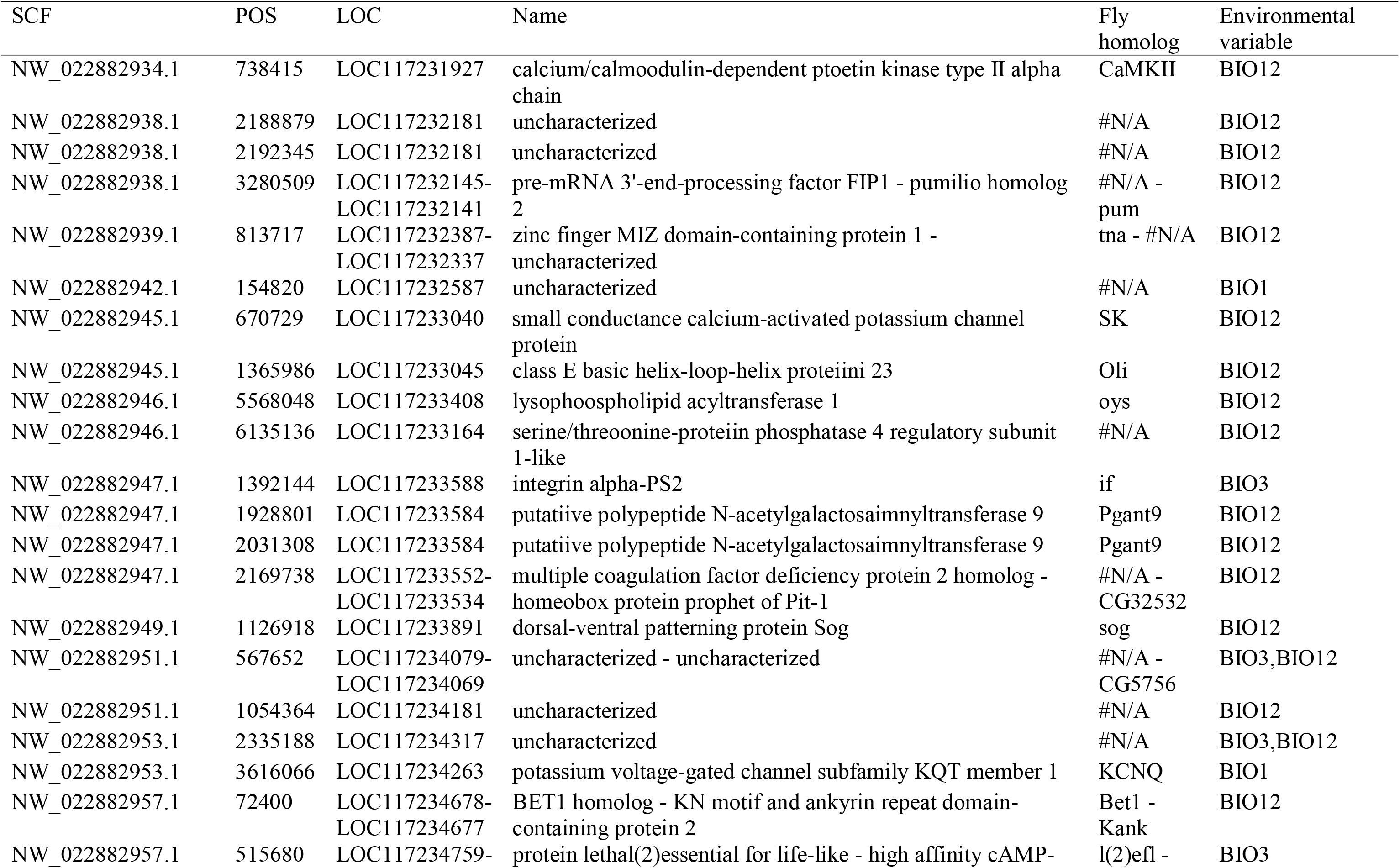
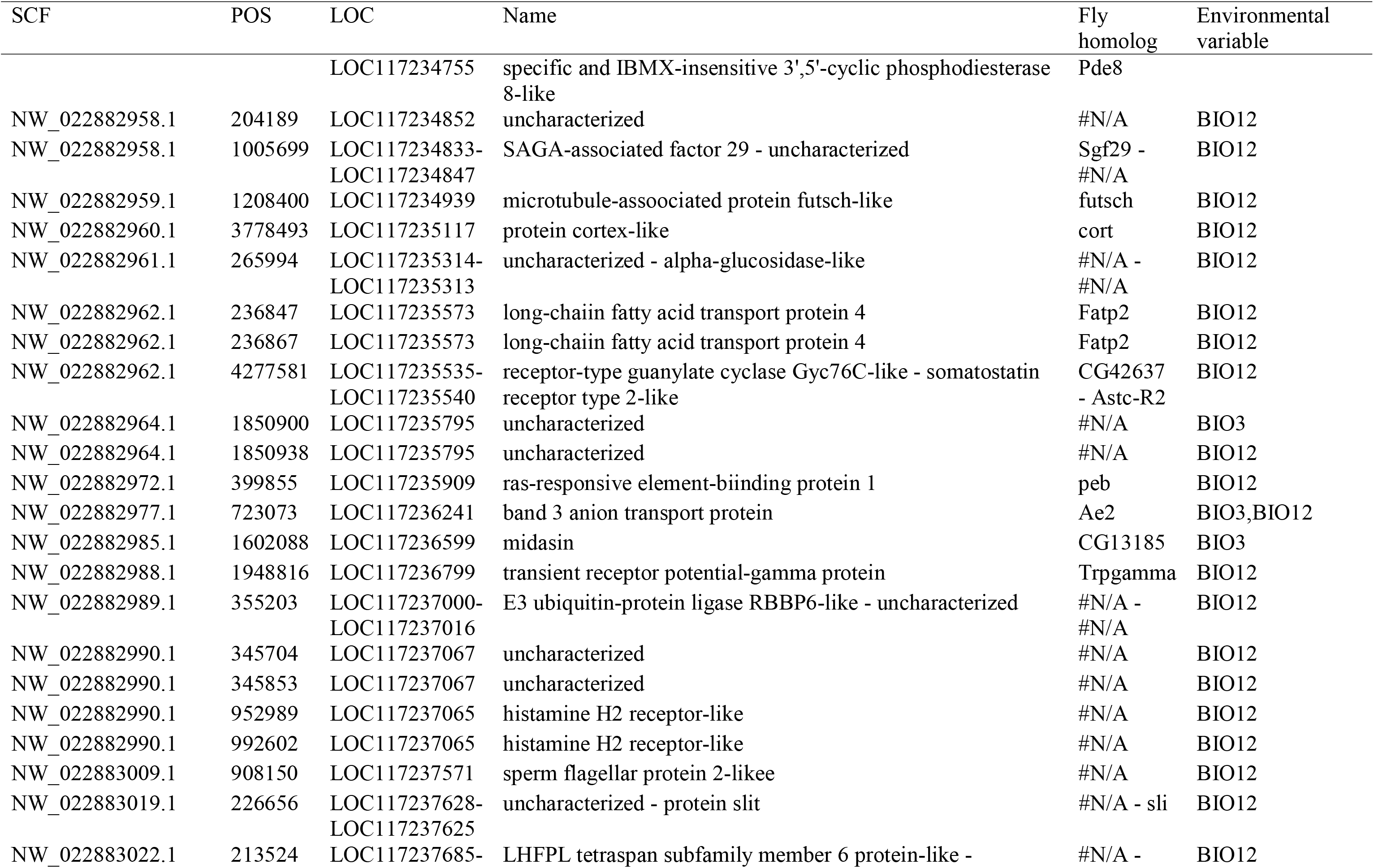
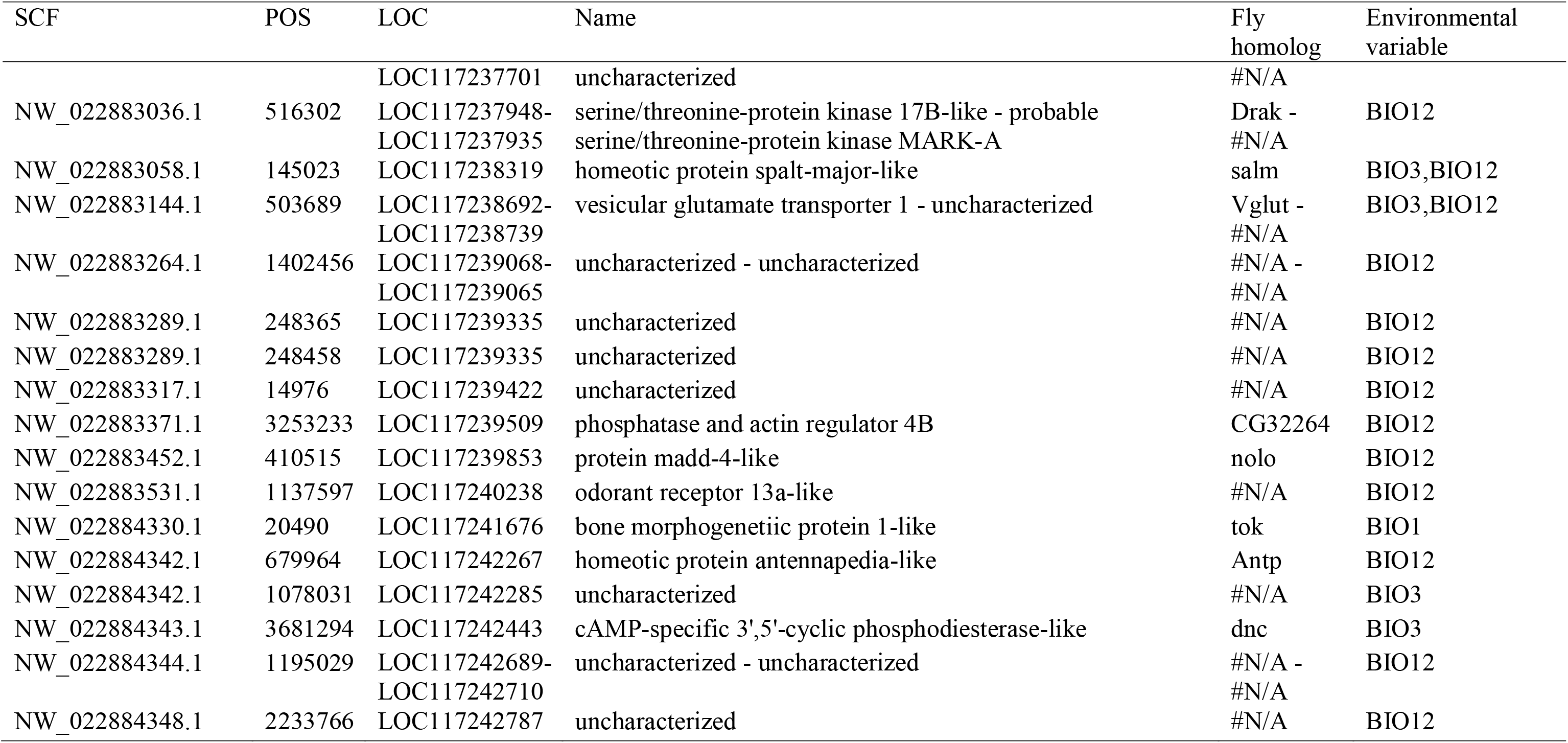
SNPs that were identified as environmentally associated outliers noting the scaffold the SNP is located on (SCF), the position of the of the SNP (POS), the NCBI identifier (LOC), the NCBI gene name (Name), the homologous gene in *D. melanogaster* if available (Fly homolog), and the environmental variable(s) associated with the SNP.

Based on the major latitudinal pattern in population structure we tested for an effect of latitude on genetic variation, and found a significant decline in π with latitude (linear regression, *F*_1,10_ = 9.465, *P* = 0.012, Fig 3a.). Despite this latitudinal trend, however, PSMC indicated similar *N*_e_ trajectories across samples that indicate a shared demographic history for genomes across the species range, consistent with the minimal overall population structure in the species (Fig 3b). Inferred population sizes were fairly stable over time, being smallest at the oldest time scales (before the last interglacial period), changing little from the Last Interglacial period to the Last Glacial Maxima aside from a modest increase just before the LGM and decline during or immediately following the LGM (Fig 3). All populations showed some degree of recent increases in *N*_e_, although the variation among genomes in the most recent time interval is likely in part driven by challenges to modeling the most recent time segments with SMC methods (Beichman et al., 2018).

**Figure 3:**
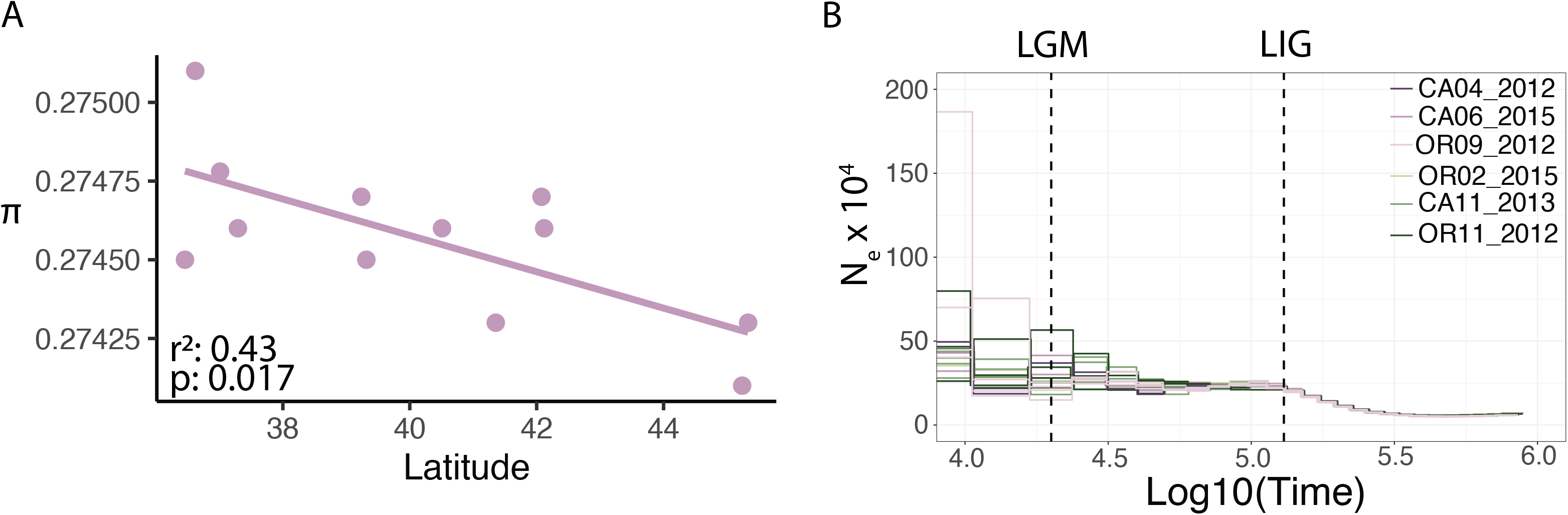
A) Demographic inference from PSMC models, marked with the approximate dates of the last glacial maximum (LGM - 20,000 years ago) and last interglacial period (LIG – 130,000 years ago) B) Linear regression between population π and latitude.

### 3.3 Environmentally associated outlier SNPs

Only a single population cluster was identified as optimal with sMNF based on the lowest cross entropy, so *k*=1 was used for LFMM2. A total of 81 loci in 72 genes were found to be significantly associated with at least one environmental variable. Significant SNPs were sparsely spread across the genome with few genes having multiple outlier SNPs and no clear peaks of association (Table 2, Fig 4). Most outlier SNPs fell within genes (60 of 81) and were principally associated with annual precipitation (n = 66, BIO12), with smaller numbers of SNPs associated with annual mean temperature (n = 4, BIO1) and isothermality (n = 16, BIO3). No outliers were associated with elevation. BIO3 and BIO12 shared 9 outlier SNPs, but BIO1 outliers were all unique.

**Figure 4:**
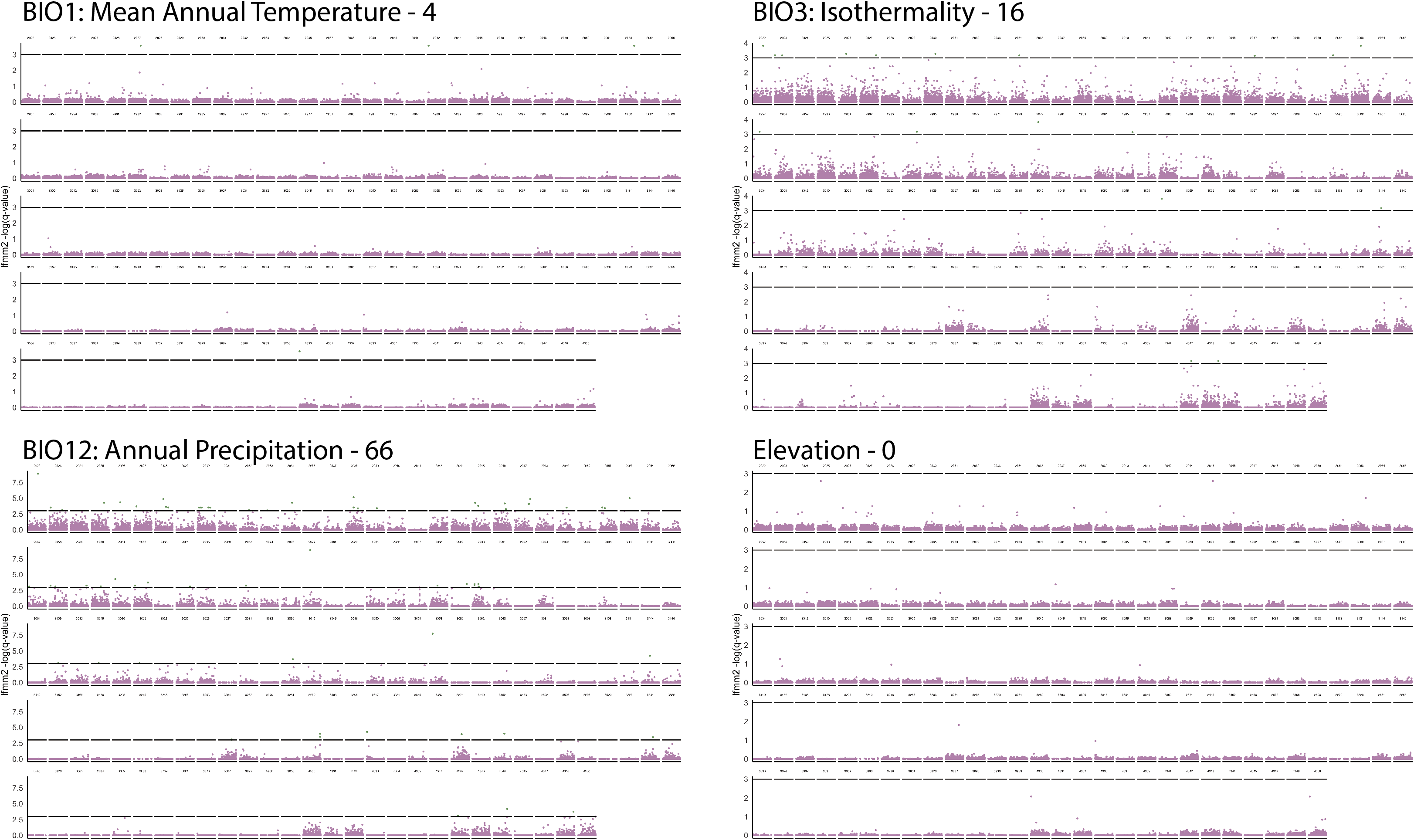
Manhattan plot of LFMM2 *q*-values for each of the four selected environmental variables (BIO1, BIO3, BIO12, and elevation). The black line represents a *q-*value threshold of 0.05, which is the threshold for significance.

Several notable genes from the LFMM2 results included LOC117231927 (*calcium/calmodulin-dependent protein kinase type II alpha chain*, homologous to *CaMKII* in *D. melanogaster*) which had the most outlier SNPs (n=3; two with BIO3 and one with BIO12). This gene has roles in neuronal growth, calcium signaling, as well as learning associated with appetite (Gillespie & Hodge, 2013). There were also several other genes with functions relating to neuronal/neuromuscular function such as LOC117231202 (*hemicentin-1*, homologous to *nrm* in *D. melanogaster*) (Kania, Han, Kim, & Bellen, 1993) and LOC117241676 (*bone morphogenetic protein 1-like*, homologous to *tok* in *D. melanogaster*) (Serpe, Ralston, Blair, & O’Connor, 2005). Interestingly, there were also several outliers in gene pairs with some form of known interaction; LOC117231348 (*potassium voltage-gated channel protein eag*, homologous to *eag* in *D. melanogaster*) is regulated by LOC117231927 (*CaMKII* in *D. melanogaster*) (Bronk et al., 2018) and LOC117233891 (*dorsal-ventral patterning protein sog*, homologous to *sog* in *D. melanogaster*) has its product cleaved by LOC117241676 (*tok* in *D. melanogaster*) (Serpe et al., 2005).

## 4 Discussion

Whole-genome resequencing of *B. vosnesenskii* across large latitude and altitude gradients in the western USA confirmed the presence of weak overall population structure and patterns of environmentally associated genomic variation that match prior results with other genetic markers (J. M. Jackson et al., 2018, 2020; Lozier et al., 2011). However, WGR data did provide some new insights, indicating that populations appear to have shared fairly concordant demographic histories (reflected by *N*_e_) over time, suggesting that the minimal population structure apparent today has persisted for thousands of years. Gene flow appears extensive across the study area and may be a likely explanation for the relatively small amount of genomic variation associated with the environmental variables and homogenous evolutionary histories reflected in genomes from different regions. Although there were relatively few and sparsely distributed environmentally associated outlier SNPs, there were some notable patterns highlighting the potential importance of ion homeostasis and neuromuscular function that is consistent with other bumble bee studies (Heraghty et al., 2022; Huml et al., 2023; Sun et al., 2020). Overall, the results suggest that *B. vosnesenskii* may be an example of a species with minimal population structure impeding adaptation or with weak selection over gene networks that will require much more extensive sampling to resolve.

Analysis of population structure suggests *Bombus vosnesenskii* is nearly but not completely panmictic, with weak isolation by distance and much of the explainable genetic variation associated with latitudinal separation, which is consistent with previous studies (Cameron et al., 2011; J. M. Jackson et al., 2018; Lozier et al., 2011). We were interested in determining whether the greater resolution afforded by WGR data would provide additional evidence for gene flow barriers across the sampled regions, but no major differences emerged compared to prior studies. Low genetic differentiation over large spatial scales, is often observed in bumble bee species in the absence of obvious physical dispersal barriers (Christmas et al., 2022; Heraghty et al., 2022; J. B. Koch, Looney, Sheppard, & Strange, 2017; Lozier et al., 2011). Although long distance dispersal of individual reproductives (gynes or drones) may be rare (Williams et al. 2022), stepping stone dispersal through suitable habitat is likely in bumble bees (Williams et al., 2022; Williams, Lobo, & Meseguer, 2018) and would contribute to weak genetic structure when species inhabit continuous geographic ranges. The isolation by distance observed here is consistent with such a model of stepping-stone population structure. *Bombus vosnesenskii* is abundant and can be found at low and high elevations throughout California and Oregon, and thus its range in these regions is composed almost entirely of contiguous suitable habitat. Not all bumble bees show this pattern, however, and data from other species in the region have higher levels of population structure. For example, *B. vancouverensis,* a species in the same subgenus (Williams, Cameron, Hines, Cederberg, & Rasmont, 2008) with a similar latitudinal range, but more narrow higher elevation niche at any given latitude (J. Koch et al., 2012), has much greater range-wide *F_ST_* based on whole genome and RADseq data (Heraghty et al., 2022; J. M. Jackson et al., 2018). Importantly, however, our sampling did not include strongly isolated populations, such as those on islands, or populations from many coastal regions, which may harbor unique genetic variation (S. Jha, 2015). Incorporating whole genome data from such populations may provide evidence of the potential for greater population structure in *B. vosnesenskii*.

PSMC historical demographic analyses were also consistent with contemporary patterns of population structure. All individuals showed highly similar historical *N_e_* trajectories, likely indicating a relatively high degree of homogeneous population structure through much of the recent past. Again, results in *B. vosnesenskii* can be compared to *B. vancouverensis*, where SMC analyses inferred markedly different bottleneck and expansion magnitudes across the species range that were linked to major climate fluctuations and consequent changes in suitable habitat areas over time that may have promoted genetic divergence (Lozier et al., 2023). For *B. vosnesenskii*, there were relatively small signatures of genomic bottlenecks or other major *N*_e_ fluctuation associated with glacial-interglacial periods, possibly suggesting that flexibility in environmental requirements may facilitate population stability and near-panmixia over time.

This would also be consistent with the minimal environmentally associated genomic variation observed here and in earlier RADseq results (Jackson et al. 2020). Despite, the minimal changes in *N*_e_ associated with changing historical conditions, there is subtle yet significant decline in nucleotide diversity with latitude, a pattern that was previously observed in microsatellite data (Lozier et al. 2011). Such patterns may provide some evidence for post-glacial northward expansion that may be shaping variation in *B. vosnesenskii*, despite the similarity in PSMC trajectories among samples, but would also be consistent with the subtle differences in the strength of very recent *N*_e_ increases observed in several of the genomes (Fig 3b).

### 4.1 Environmentally associated outliers

The lack of major population structure and parallel *N*_e_ histories across the *B. vosnesenskii* range is mirrored by a lack of clear influence of environmental variables on overall population structure or on allele frequencies at individual loci that might be produced by local adaptation.

Interestingly, most of the detected outliers were associated with precipitation, with smaller numbers of loci associated with other variables, and none with elevation. Although, the outlier loci were sparsely spread throughout the genome (Table 2, Fig 4), there were several interesting cases where outlier-containing genes interacted, possibly suggesting the potential for weak selection on members of gene networks. This is supported by several instances where outlier genes have known regulatory interactions (e.g., LOC117231927 and LOC117231348 or LOC117233891 and LOC117241676). The overall effect of selection across networks can be evaluated by looking at the roles of the individual genes with detected outliers, such as the number of outlier-containing genes related to neuromuscular function.

Given the broad latitudinal and elevational range sampled for this study, the scarcity of outlier SNPs associated with thermal variables (BIO1 and BIO3) in these whole genomes was somewhat unexpected but does match the less dense RADseq SNP set results from Jackson et al. (2020) that included even more individuals and populations. One possible explanation is that *B. vosnesenskii* is a noted elevation generalist (J. Koch et al., 2012; Lozier et al., 2021; Thorp, Horning Jr., & Dunning, 1983), which, combined with high gene flow, may not produce strong or consistent signatures adaptations for elevation or temperature stressors like isothermality across the species range. The scarcity of outliers tied to elevation or temperature is similar to the absence of correlations between relevant functional morphological traits (e.g., body size and wing loading) with temperature or elevation observed in this species (Lozier et al., 2021).

However, a lack of genome regions that show clear associations with temperature is still surprising given recent work on thermal tolerance physiology in *B. vosnesenskii* (Pimsler et al 2020) that found populations from cold, high elevation habitats had significantly lower critical thermal minima (CT_min_) compared to populations from warmer localities (Pimsler et al., 2020). It is possible that selection is weak on individual genes associated with a complex trait like thermal tolerance (Yang, Crossley, Schrader, & Dubovskiy, 2022), that different genes may be targeted by selection in different regions (Yeaman, 2022), or that another non sequence-based mechanism (e.g., epigenetic) could facilitate variation in cold tolerance among populations (Mccaw, Stevenson, & Lancaster, 2020). Detecting selection under the high levels of gene flow that seems prevalent in *B. vosnesenskii* will also be challenging, and more independent studies, additional sampling, and experimentation will be needed to fully understand any mechanisms of genetic adaptation in this species.

In addition to the general challenges of sample size when selection is weak on individual genes, there are several other important considerations when evaluating adaptive potential of individual outliers. First, environmental variation is spatially correlated, so models need to consider species demography (Hoban et al., 2016). Here, population structure is weak and the statistical method employed (LFMM2) accounts for population structure, which may limit major effects here, but at the same time LFMM2 is a relatively conservative method (Luo, Tang, Schoville, & Zhu, 2021), which might contribute to reduced power to detect outliers with weak environmental associations with our modest sample size (Ahrens et al., 2018). Second, correlation between environmental variables can complicate linking environmentally associated loci to driving pressures. For instance, annual precipitation, which had the most outlier loci, could be associated with a number of different potential pressures such as desiccation tolerance, changes in biotic interactions (e.g., flowering phenology), or a number of other factors (see (Chown, Sørensen, & Terblanche, 2011) for a more thorough discussion). The presence of outliers that are common to multiple environmental variables (e.g., BIO12 and BIO3) highlights this challenge, and indicates that even a strong signal of possible selection could be associated with multiple (even unsampled) variables.

The outlier signatures in *B. vosnesesnskii* can be summarized by once again comparing to *B. vancouverensis* from similar geographic regions (Heraghty et al., 2022; J. M. Jackson et al., 2020). From both whole genome resequencing and RADseq SNPs, *B. vancouverensis* has much stronger signals of environmental association across its genome, with several large peaks of genomic divergence in genes with putatively relevant functions (Heraghty et al., 2022). Thus, in comparison to *B. vancouverensis*, *B. vosnesenskii* has less substantial population structure, a lack of population-specific demographic histories from SMC analyses, and much sparser evidence for local adaptation. That said, our results here do reveal a few parallels between these species with respect to the functions of genes with identified outlier SNPs. For instance, the gene LOC117231202 (*hemicentin-1*) was identified as containing outlier SNPs in both species. This gene reflects the shared trend towards genes involved in neuromuscular function (Kania et al., 1993) in outlier sets for both species, which has also been observed in outlier analysis of other bumble bees (Huml et al., 2023). Although elevation and thermal variables were not as associated with outliers in *B. vosnesenskii*, given correlations among environmental variables discussed above, such processes could nonetheless reflect shared selection pressures on traits such as thermal adaptation and flight. Bumble bee flight is crucial for foraging, dispersal, and overall colony fitness (Mola et al., 2020a), while the capacity for thermogenesis by shivering their flight muscles is well-recognized in *Bombus* (Heinrich 1975). These overlaps may suggest there may be some common tools important for bumble bees that occupy spatial-environmental heterogeneity across their ranges, even if such species exhibit different intensities in evidence for selection, which would be well worth additional research.

### 4.2 Broader Implications

As one of the most common bees in the western U.S.A., *B. vosnesenskii* is an important pollinator in natural ecosystems as well as agricultural systems in California and Oregon (Fisher, Watrous, Williams, Richardson, & Woodard, 2022; Greenleaf & Kremen, 2006). Thus, ensuring the health of *B. vosnesenskii* populations in a changing environment is of great interest. Our results contribute to the current understanding of how *B. vosnesenskii* may fare under climate change. Contemporary analysis of *B. vosnesenskii* suggests that this species has remained stable over recent decades (Cameron et al., 2011). The historical demographic inferences from SMC modeling suggest that populations sizes across our study region have likewise been minimally affected by the change in environmental conditions from the LIG to the LGM (Fig 3). During the LIG, conditions were warmer than present by ∼1.2 °C relative to the 1998-2016 average (Lüning & Vahrenholt, 2017) whereas LGM conditions were colder by ∼6 °C (Von Deimling, Ganopolski, Held, & Rahmstorf, 2006). The species’ success under warmer conditions is particularly encouraging since it may indicate stability in future climates (IPCC 2021). Recent work suggests that higher temperatures may even be beneficial to *B. vosnesenskii* and a possible factor in range expansions into British Columbia (Fraser et al., 2012; H. M. Jackson et al., 2022), with *B. vosnesenskii* representing a possible “climate winner”, at least to date (Jackson et al. 2022). In addition to temperature, precipitation patterns will also shift in future climate change (IPCC 2021). Our results suggest that annual precipitation is associated with most genomic selection signatures, and prior results found precipitation to be a key predictor of *B. vosnesenskii* range limits in species distribution models (Jackson et al. 2018). Although recent work has indicated that changes in precipitation are less likely to have an impact on future distributions of North American bumble bees, there are some exceptions (H. M. Jackson et al., 2022), and our genomic data indicate that water limitation may represent an important consideration under future climates that may warrant further study.

In summary, we have confirmed previous findings indicating there is relatively little population structure in *B. vosnesenskii* (J. M. Jackson et al., 2018) and identified relatively few environmental associated outliers which could underlie adaptation in this species. The lack of outliers could be associated with a gene swamping effect resulting from the high observed gene flow (Lenormand, 2002), although contradicts data that suggests biogeographic variation in key thermal performance metrics (Pimsler et al. 2021). For the environmentally associated outliers that were identified, future work should be conducted to evaluate their potential role in local adaptation. Additionally, future work should be aimed at teasing apart the environmental variation that is captured by precipitation (BIO12) and how that might impact adaptation and future species distributions, given changes in precipitation patterns forecasted under climate change. Finally, more work is needed to examine other avenues for adaptation in this species, such as epigenetic modifications which may explain observed differences in cold tolerance across the range.

## Acknowledgements

We thank the university of Alabama College of Arts and Sciences and the National Science Foundation (DEB-1457645 and URoL 1921585 to J.D.L.) for support related to this project. We also thank the other members of the Lozier lab for useful feedback and discussion.

## Data Accessibility

Raw sequencing data is available on NCBI SRA (Bioproject PRJNA909611; accession numbers SAMN32298940- SAMN32298956, SAMN32230877 - SAMN32230970 SAMN32298940).

Variant data for SNPs as well as scripts used in data filtering and analysis are available on FigShare.

## Author Contributions

S.D.H. processes samples, performed statistical analysis, wrote the manuscript, and edited the manuscript. J.M.J. collected samples and helped to process samples. J.D.L. designed the study, obtained funding, conducted field work, helped with statistical analysis, and edited manuscript.

